# Ontogenesis of the molecular response to sleep loss

**DOI:** 10.1101/2023.01.16.524266

**Authors:** Christine M. Muheim, Kaitlyn Ford, Elizabeth Medina, Kristan Singletary, Lucia Peixoto, Marcos G. Frank

## Abstract

Sleep deprivation (SD) results in profound cellular and molecular changes in the adult mammalian brain. Some of these changes may result in, or aggravate, brain disease. However, little is known about how SD impacts gene expression in developing animals. We examined the transcriptional response in the prefrontal cortex (PFC) to SD across postnatal development in male mice. We used RNA sequencing to identify functional gene categories that were specifically impacted by SD. We find that SD has dramatically different effects on PFC genes depending on developmental age. Gene expression differences after SD fall into 3 categories: present at all ages (conserved), present when mature sleep homeostasis is first emerging, and those unique to certain ages in adults. Developmentally conserved gene expression was limited to a few functional categories, including Wnt-signaling which suggests that this pathway is a core mechanism regulated by sleep. In younger ages, genes primarily related to growth and development are affected while changes in genes related to metabolism are specific to the effect of SD in adults.

## 1. Introduction

Studies in adult animals show that sleep deprivation (SD) results in wide-spread changes in brain gene expression (Cirelli et al., 2004; Gerstner et al., 2016; Mackiewicz et al., 2007; Maret et al., 2007; Naidoo et al., 2005; Terao et al., 2006). These studies have provided key insights into mechanisms governing the deleterious effects of sleep loss as well as those that mediate sleep drive. For example, SD down-regulates transcriptional factors important for mRNA translation, which in part may explain some neurological deficits associated with abnormal sleep (Gerstner et al., 2016). SD also up-regulates many immediate early genes, including those hypothesized to play essential roles in sleep drive and sleep function (*e.g*. Homer1a (Cirelli et al., 2004; Maret et al., 2007)).

Far less is known about molecular changes that accompany sleep loss in developing animals. This is an important, unexplored area in sleep biology because sleep in early life may play critical roles in brain development, plasticity, and neurodevelopmental disorders such as autism (Blumberg et al., 2020; Frank, 2011a; Medina et al., 2022). In addition, developing animals do not respond to sleep loss the same way as adults, indicating that the accumulation and discharge of sleep drive undergoes important transformations during the postnatal period (Davis et al., 1999). This has been best described in rodents, where until a certain postnatal age, SD fails to elicit the normal adult compensatory increase in non-rapid-eye-movement (NREM) electroencephalographic (EEG) slow-wave activity (Frank et al., 1998a). These latter findings suggest that molecular changes triggered by SD may also change across postnatal development.

We explored this possibility using cortical RNA sequencing in male mice at time points that span ages when the response to SD is immature and then adult-like. This was combined with analytical techniques that increase power and reproducibility of results in genomic data (i.e., *removal of unwanted variance*: RUV). We find that at ages that correspond to an immature response to SD, cortical gene expression is dramatically different than what is observed in adult animals. We also find that there is a small, but conserved set of genes that change in similar ways after SD at all ages; and that Wnt-signaling is affected by sleep loss at all ages but particularly so at postnatal day 24. These findings suggest that the cost of wakefulness changes across development and there is a conserved function of sleep that emerges very early in development.

## 2. Materials and Methods

### 2.1 Experimental model and subject details

Wild-type (WT) C57BL6/J male mice were used. Mice were housed in standard cages on a 12:12 h light:dark cycle with food and water *ad libitum*. Adult mice (10-12 weeks old, hereafter called P90) were individually housed 7 days before the experiment. P24 mice were weaned at P18 and P30 were weaned at P24, respectively, then housed separately. P16 mice were separated from the dam at ZT0 on the day of the experiment and housed with a littermate in a warmed cage (gradient 21-29°C) enriched with housing material from the breeding cage. Animals were divided into 2 groups; sleep deprived (SD), and home cage controls (HC), n=5-8 independent animals per group. HC control mice were left undisturbed and were euthanized either 3h (P16, P24, P30) or 5h (P90) after light onset (ZT3, ZT5). These SD times were selected based on previous reports indicating that rodents <P30 cannot stay continuously awake for more than 3-4 hours, and SDs of these durations are sufficient to induce a homeostatic increase in sleep time or intensity (Alfoldi et al., 1990; Blumberg et al., 2004; Frank et al., 1998a; Gvilia et al., 2017, 2011; Medina et al., 2022). All experimental procedures were approved by the Institutional Animal Care and Use Committee of Washington State University and conducted in accordance with National Research Council guidelines and regulations for experiments in live animals.

SD for the P24, P30, and P90 was performed as described previously (Muheim et al., 2021). Briefly, single housed animals were kept awake with gentle handing (e.g. stroking with a soft brush to keep mice moving and exploring). These techniques have been previously shown to keep developing rodents awake >90% of the time during the SD (Frank et al., 1998b; Medina et al., 2022). For the P16 group, animals were kept with a sibling in a warmed cage and initially only touched with a soft brush. When the soft brush did not successfully keep them awake anymore, mice were gently kept awake by passing them slowly from one hand to the other.

### 2.2 Tissue collection and RNA isolation, library preparation, and sequencing

RNA isolation was performed as previously described (Ingiosi et al., 2019). Briefly, mice were euthanized upon completion of SD by cervical dislocation without prior sedation to avoid the confounding effects of anesthesia on gene expression. PFC was dissected and RNA was extracted using the Qiagen RNAeasy kit (Qiagen). RNA-seq library preparation, quantification, and sequencing was performed at the WSU Spokane Genomics core. Library quantification was done using the KAPA quantification kit (Kapa Biosystems, Wilmington, MA). Sequencing was accomplished using HiSeq 2500 technology (Illumina SBS kit v4) at an average depth of 44 million 100bp pair-end reads per sample.

### 2.3 RNA-seq data analysis

Transcript quantification was done with Salmon (v1.8) (Patro et al., 2017) using the GENCODE reference genome and transcriptome (release M28). To improve the accuracy of quantification estimates the salmon index incorporates decoys (Patro et al., 2017). The R/Bioconductor package tximeta (v1.14)(Love et al., 2020) was used to create a gene level matrix of transcript abundance for all samples. Gene counts were filtered to require at least 10 reads over 5 samples to avoid zero inflation. Removal of unwanted variation (RUVs) normalization was performed using RUVseq (v1.30) (Peixoto et al., 2015; Risso et al., 2014) as in our previous work (Ingiosi et al., 2019). Briefly, RUVs (k=3) was applied after samples were grouped based on age and treatment using both negative controls genes and samples (biological replicates). To obtain control genes, in our previous work, four different genome-wide datasets from four different brain tissue sources and three different laboratories were integrated to define differentially expressed genes after 5-6 hours of SD in adult male mice (Gerstner et al., 2016). Negative control genes were defined as those unaffected by SD (uncorrected p-value >0.8, supplementary Table 1) independent of tissue and lab. A list of 572 positive control genes (FDR <0.05, supplementary Table 1) from the same study was also used to evaluate the reproducibility of the differential expression analysis in adult animals.

Differential expression analysis was run using edgeR (v3.38) with a factorial design that included age and treatment (HC and SD) at an FDR <0.05. To find sets of genes that are either unique to each age or shared between ages, we used Venn diagrams to define subsets of interest for biological function analysis.

Functional annotation analysis of gene sets of interest was done using the Database for Annotation, Visualization, and Integrated Discovery v2021 (DAVID, https://david.ncifcrf.gov (Huang et al., 2009a, 2009b)). The following databases of functional information were used: Uniprot keywords for Molecular Function (MF), and Biological Process (BP, https://www.uniprot.org), and KEGG pathways (PW, https://www.genome.jp/kegg/pathway.html). Enrichment was determined relative to all genes expressed in the mouse PFC based on our data and with at least three genes per function or pathway. Clustering of functions or pathways was performed as previously described (Huang et al., 2009c) using a similarity threshold of 0.2 and an enrichment p-value of <0.05. Log2 fold change of genes in functions or pathways with a fold enrichment >4 were generated using heatmap.2 (R package ggplot2_3.3.6). Clustering of genes for the heatmaps were determined using Euclidean Distance with the Ward D2 agglomeration method.

### 2.4 Data availability

Sequencing data have been deposited in the Gene Expression Omnibus Database (GEO) under accession number GSE211301. Of the 16 adult WT PFC samples, 10 were obtained from GSE113754 (5 SD, 5 HC). Code used for the analysis and visualization is available on Github under: https://github.com/PeixotoLab/MolecularSleepOntogenesis.

## 3. Results

Our main findings can be summarized as follows. First, the number of differentially expressed genes caused by SD increases with developmental age. Second, these genes fall into three main classes: 1) age independent, 2) specific to ages with adult-like sleep homeostasis and 3) age specific. Genes in class 1 regulate transcription, genes in class 2 are involved in cellular stress response, proteostasis, and circadian rhythms. Genes in class 3 are related to development and growth for juvenile (P24) and metabolism for adult (P90) mice. Only one signaling pathway was enriched in genes differentially expressed regardless of age: Wnt-signaling, with the most pronounced effects at P24. These results are presented in greater detail in the following sections.

### 3.1 SD-induced transcription changes with age

As we have previously shown (Gerstner et al., 2016; Ingiosi et al., 2019; Peixoto et al., 2015), RUV normalization is necessary to properly detect the effects of the variables of interest (age and SD). Fig.1A shows principal component analysis (PCA) of the data displaying a clear grouping of biological replicates per treatment group, while traditional normalization methods that rely on correcting sequencing depth cannot properly capture the treatments (Fig.S1A). Further, the dynamic expression range after RUV normalization is less variable within groups and more balanced across groups (Fig.1B, S1B) and the number of positive control genes in adult mice is larger compared to sequencing depth normalization (84.4% vs 71.6% Fig.1F, S1F). Therefore, RUV normalization is essential to both the power and reproducibility of downstream analyses. Following proper normalization, our differential expression analysis shows that increased age was accompanied by a step wise increase in expression change (Fig.1C-F). Although only 85 genes are differentially expressed after SD at P16, that number increases to 3601 at P24, 2423 at P30, and 6370 at P90 (supplementary Table 2). In general, we see more downregulated genes than upregulated genes in all groups.

**Figure 1.**
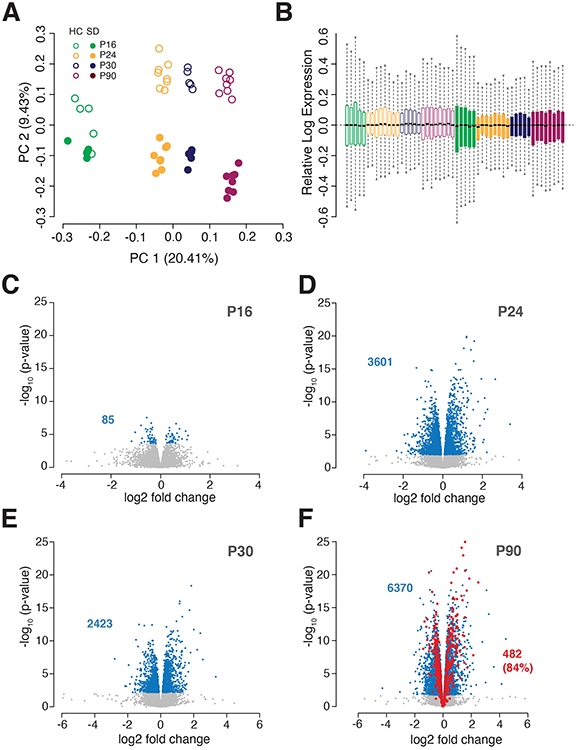
The number of cortical genes influenced by SD increases with age. **A)** Principal component analysis of RUVs normalized sequencing reads showed an increased effect of sleep-deprivation with increased age. Developmental age contributed the most variability (PC1, 20.41%) while SD contributed only 9.43% (PC2) to the overall variability in the dataset. n=5-8 per age group and condition, k=3, open circles: home cage, closed circles: SD. **B)** Relative Log expression for each sample and condition using k=3 factors for RUVs. Color code as in A. **C-F)** Volcano plots of normalized dataset per age group, **C)** P16 **D)** P24 **E)** P30 **F)** P90. Each gray dot represents an expressed gene, blue dots are differentially expressed genes after SD with the absolute gene number in blue for each volcano plot on the left. For the P90 group, red dots represent control genes with known response to SD (supplementary Table 1). 84.4%of these genes were found as differentially expressed after SD with the chosen RUVs normalization, k=3, FDR <0.05.

Next, we determined which differentially expressed genes after SD are conserved across ages. Only 59 genes are differentially expressed after SD at all ages (29 up, 30 down, Fig.2, dark green outline, supplementary Table 3). A larger group of 1563 genes is shared in the response to SD between P24, P30, and P90 (689 up, 874 down, Fig.2, yellow outline, supplementary Table 3).

**Figure 2.**
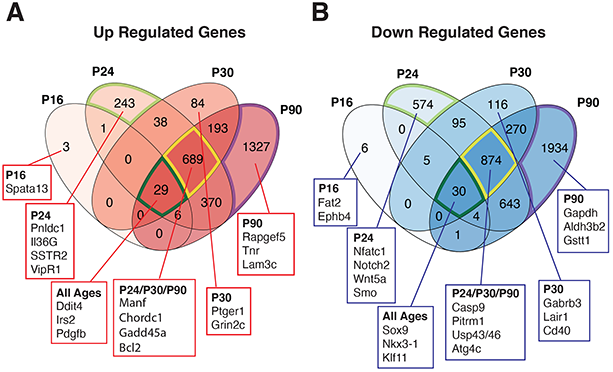
Venn diagram of **A)** up and **B)** down regulated genes after SD in each age group with an FDR of <0.05. Selected genes in the boxes are given as examples for each category analyzed further. Outlined in dark green, yellow, lime green or purple are the subsets of genes further analyzed. Dark green: genes changed after SD in all ages, yellow: genes changed only in P24, P30, and P90, lime green: genes changed exclusively in P24, purple: genes changed exclusively in P90.

To better understand the biological implications of the differentially expressed genes after SD across ages, we ran a functional enrichment analysis to determine which pathways or biological functions are over-represented in specific subsets of genes relative to what is expected by chance. To do so we defined the following subsets of interest: 1) differentially expressed genes after SD conserved across ages 2) differentially expressed genes after SD shared between P24, P30, and P90 and 3) differentially expressed genes after SD unique to each age group.

### 3.2 The conserved response to SD regardless of age involves downregulation of transcription and Wnt-signaling

Genes regulated by SD at all ages were primarily involved in transcriptional regulation. From the very small set of conserved genes (Fig.2, dark green outline, 29 up, 30 down, supplementary Table 3) we could identify 7 enriched functions or pathways, of which 6 involve downregulated genes and only 1 upregulated gene (Fig.3A, supplementary Table 4). All downregulated 6 functions are related based on the clustering results. This suggests that sleep loss per se reduces overall transcriptional activity reflected by less DNA binding or the reduced abundance of transcriptional activators such as *Sox9*, *Nkx3-1,* or *Klf11*. Wnt-signaling is the only pathway enriched, based on the downregulation of *Wnt9a* (Lyons et al., 2020), *Fzd10,* and *Nfat3c*. For upregulated genes, we only found enrichment of genes that are targeted by microRNAs in cancer (*Ddit4*, *Irs2*, *Pdgfb,* and *Cdkn1a*). When comparing the direction of change across ages for the genes in the most enriched functions (>4-fold), the change in expression increases with age (Fig. 3B).

**Figure 3.**
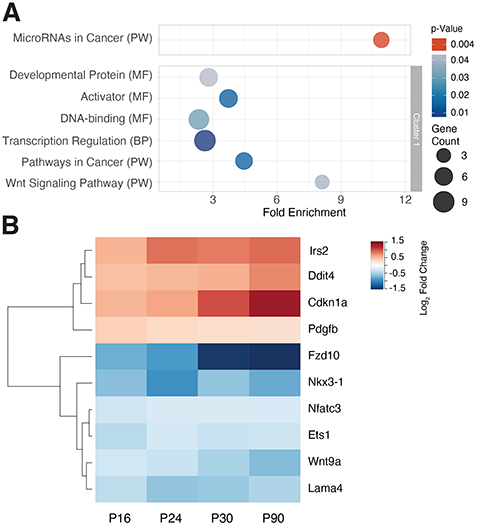
Genes affected by SD at all ages target transcription. **A)** Bubbles show functional enrichment analysis of up (red) and down (blue) regulated genes. Bubble size represents number of genes per functional category. Modified Fisher’s exact p-value<0.05, minimum of 3 genes for corresponding functions or pathways. P-values are represented as a color gradient. Gray box indicates functions within a cluster with an enrichment score of 1.65 based on a similarity threshold 0.2 and Modified Fisher’s exact p-Value<0.05). No significant cluster was detected for the upregulated genes. MF, Uniprot keywords Molecular Function. BP, Uniprot keywords Biological Process. PW, KEGG pathways. **B)** Heatmap of log2 fold change across age of genes enriched in functional categories with >4 fold enrichment from A (MicroRNAs in Cancer, Wnt-signaling Pathways, Pathways in Cancer). Red colors indicate increase in expression compared to home cage control, blue colors indicate decrease. Dendrogram on the left represents gene clusters based on Euclidean distance and Ward 2D agglomeration of log2 fold changes.

### 3.3 Genes regulated by SD at P24 and older are involved in protein stability and circadian rhythms

The largest subset of shared genes induced by SD are between P24, P30, and P90 (1563, 689 up and 874 down, Fig.2, yellow outline, supplementary Table 3). These genes are of special interest because they could indicate the genetic origin of mature sleep homeostasis. The response to SD as seen in adult animals, emerges at P24 and intensifies with age until adulthood, but is absent in earlier ages (Frank et al., 1998a; Medina et al., 2022). Genes differentially expressed at these ages but not earlier might therefore be related to the regulation of sleep homeostasis. In our study, functional enrichment followed by clustering analysis revealed that genes related to proteostasis (such as the unfolded protein response), immune responses, and biological rhythms were significantly enriched. We found 24 upregulated functions, some of which are unrelated to each other and some of which constitute two clear clusters (of 4 and 13 functions respectively, Fig.4). The first cluster, involves stress response (e.g. *Hsp5a* (Naidoo et al., 2005), *Bcl2* (Montes-Rodríguez et al., 2009) or *Xbp1* (Brown et al., 2014)), while the second cluster involves pathways regulating immune function and apoptosis (e.g., *Nkfbia* (Giannos et al., 2022) *or Sgk1* (Giannos et al., 2022; Mongrain et al., 2010)). Despite a large group of downregulated genes (874, Fig.2 yellow outline, supplementary Table 3), few biological functions or pathways were enriched, suggesting a generalized non-specific downregulation of transcription by SD from P24 onwards.

**Figure 4.**
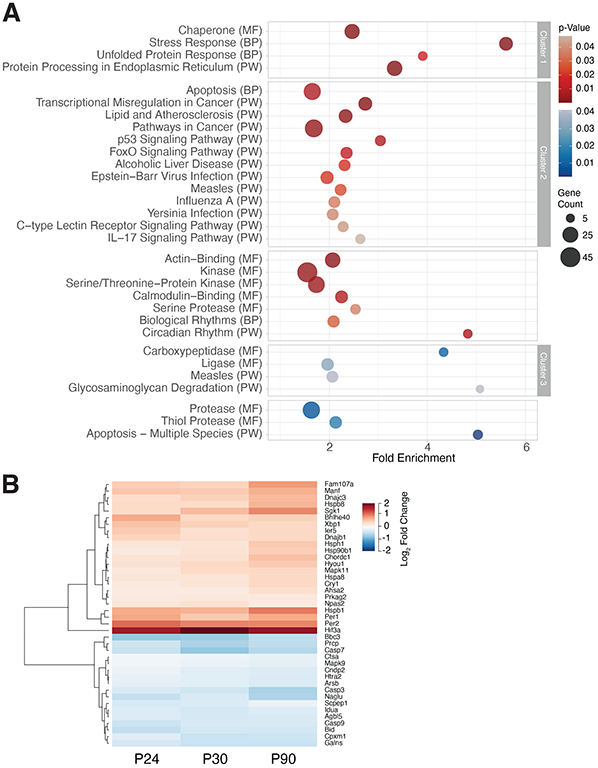
Genes affected by SD in P24 and older animals mainly affect protein regulation and circadian rhythms. **A)** Bubbles show functional enriched terms of up (red) and down (blue) regulated genes. Bubble size represents number of genes per term. Modified Fisher’s exact p-value<0.05, minimum of 3 genes for corresponding functions or pathways. P-values are represented as a color gradient. Gray boxes outline functions for each cluster based on a similarity threshold of 0.2 and a Modified Fisher’s exact p-value<0.05. Enrichment scores for each cluster; cluster-1: 4.72, cluster-2: 1.93, cluster-3: 2.17. MF, Uniprot keywords Molecular Function. BP, Uniprot keywords Biological Process. PW, KEGG pathways. **B)** Heatmap of log2 fold change across age of genes enriched in terms with >4 fold enrichment from A (Stress Response, Circadian Rhythm, Apoptosis - multiple species, Glycosaminoglycan degradation, Carboxypeptidase). Red colors indicate increase in expression compared to home cage control, blue colors indicate a decrease. Dendrogram on the left represents gene clusters based on Euclidean distance and Ward 2D agglomeration of log2 fold changes.

### 3.4 Age specific changes after SD in P24 mice are related to growth and development while changes in P90 mice are generally related to metabolism

Subsequently we focused on gene expression differences that are unique to each age. The youngest group, P16, had only 9 genes in this category with no specific functions associated with them (supplementary Table 3). P24 animals had 243 upregulated and 574 downregulated genes (Fig.2, lime green outline, supplementary Table 3) that grouped into 11 enriched biological functions or categories (3 up and 8 down, Figure 5). These include immunity and neuroactive ligand receptor interaction (e.g., *SSTR2*, *Vipr1*, *Drd1*, *Mchr1,* or *Galr1*) for upregulated genes. Functional categories over-represented among downregulated genes exclusive to P24 belong to growth factor binding, ligand-gated ion channels, tyrosine metabolism, and Wnt-signaling among other categories. Although a small subset of Wnt-signaling pathway genes are downregulated by SD regardless of age, at P24 there is a notable increase of the effect of SD towards the downregulation of this pathway (Fig.6). Genes that were uniquely affected by SD at P30 are not enriched in any biological pathway or function at our established statistical cutoff.

**Figure 5.**
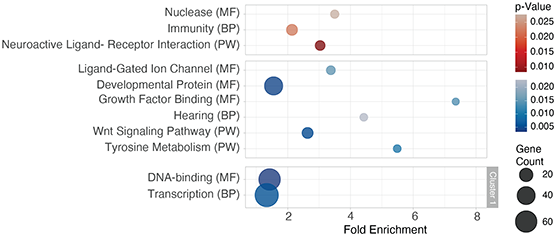
Functional enrichment analysis for differentially expressed genes after SD exclusively in P24 mice are related to development and neuronal activity. Bubble size represents number of genes per function, for up (red) and down (blue) regulated genes, Modified Fisher’s exact p-value<0.05, minimum of 3 genes for corresponding functions or pathways. P-values are represented as a color gradient. Gray box outlines enriched functions per cluster based on a similarity threshold of 0.2 and a Modified Fisher’s exact p-value<0.05, enrichment score 2.31. MF, Uniprot keywords Molecular Function. BP, Uniprot keywords Biological Process. PW, KEGG pathways.

In P90 mice, where the largest group of differentially expressed genes were observed (1327 up, 1934 down, Fig.2, purple outline, supplementary Table 3), we found 22 enriched functions with one large cluster for the upregulated group and 30 enriched functions with 5 clusters for the downregulated group (Fig.7, supplementary Table 4). Genes specifically affected by SD in P90 were largely related to cell adhesion and polarity (upregulated, e.g. *Epha2*, *Tnr,* or *Lama1*) and metabolism (downregulated, e.g. *Nos2* (Chen et al., 2003), *Gapdh,* or *Aldh3b2*, supplementary Table 4).

## 4. Discussion

This is the first study investigating the effects of SD on cortical RNA expression across postnatal development. Earlier reports (Bellesi et al., 2013; Cirelli et al., 2004; Cirelli and Tononi, 2000; Mackiewicz et al., 2007; Maret et al., 2007; Noya et al., 2019; Terao et al., 2006) used adult animals to measure the effects of SD on gene expression. We separated age specific effects from global, age-independent effects of SD and therefore provide a more detailed view of the changes SD induces on the transcriptional level. We further discuss some of these findings in more detail below.

### 4.1 Conserved gene expression after SD

We find a very small set of differentially expressed genes after SD present at all developmental ages. Among them, the Wnt-signaling pathway is significantly enriched. In developing and adult mice, Wnt-signaling is involved in a wide variety of essential functions, such as brain cell differentiation, migration, synaptic activity/plasticity and cell survival (Nusse and Clevers, 2017). However, this is the first time that SD has been shown to affect components of this essential signaling pathway. Wnt-signalling has a precise spatio-temporal organization defined by the combination of the wnt ligands and frizzled receptors (Loh et al., 2016; Willert and Nusse, 2012). The specific downregulation of *Wnt9a* (also called *Wnt14* (Qian et al., 2003)), *Fdz10,* and *Nfatc* suggest a key function for these elements in regulating neuronal activity during prolonged wakefulness. It is unclear what specific role *Wnt9a* or *Fdz10* might play in neuronal maintenance across age. It has been shown that *Wnt9a* is reduced in the aging brain (Hofmann et al., 2014) and recent findings link aberrant Wnt-signaling to neurodegenerative disorders such as Alzheimer’s disease (Palomer et al., 2019). This suggests that a core and developmentally conserved function of sleep is the regulation of neuroprotective mechanisms.

### 4.2 Cortical gene expression and the emergence of (mature) sleep homeostasis

Rodents younger than ≈ P24 do not show compensatory changes to SD based on NREM sleep EEG activity (Frank et al., 1998a; Frank and Heller, 1997a, 1997b) (the canonical measure of mammalian sleep homeostasis (Borbély, 1982; Brunner et al., 1993)). For example, in Long-Evans rats, animals younger than P24 only respond to SD with changes in sleep time or duration. Changes in NREM sleep SWA first appear around P24 (the precise timing is strain dependent (Gvilia et al., 2017, 2011)). Sleep homeostasis in developing mice has not been studied in similar detail, but in other measures (e.g. changes in sleep time and architecture) they follow similar developmental patterns as rats (Daszuta et al., 1983; Daszuta and Gambarelli, 1985; Rensing et al., 2018). The underlying reasons for this change in sleep homeostasis are unknown but may be revealed by molecular analyses as performed here. For example, Serum and Glucocorticoid Induced Kinase 1 (*SGK1*) is first up regulated by SD at P24 and is one of the few genes consistently upregulated (across studies) by SD in adult rodents (Giannos et al., 2022). *SGK1* is expressed in neurons and astrocytes (Miyata et al., 2011; Slezak et al., 2013; Wärntges et al., 2002) and plays an important role in cellular stress response, cell survival, and excitability (Wärntges et al., 2002).

Interestingly, we also observe an upregulation of other genes at P24 onwards related to cellular stress such as the unfolded protein response (UPR) or chaperones; pathways shown to be upregulated by SD in adult rodents (Aboufares El Alaoui et al., 2023; Cirelli et al., 2004; Naidoo et al., 2005). We also find that the canonical clock genes *Per1/Per2* and *Cry1* are differentially expressed from P24 onwards. In adult rodents, circadian clock genes are similarly regulated by SD (Curie et al., 2014; Franken et al., 2007; Mongrain et al., 2010; Wisor et al., 2008, 2002) and interact with damaged DNA (Özgür and Sancar, 2003). Similar mechanisms may occur at parallel developmental periods in humans, as around 2-3y of age there is a shift from neuronal organization towards repair mechanisms (Cao et al., 2020).

### 4.2 SD induced changes unique to P24: a window of vulnerability?

Our findings suggest that the rodent cortex is particularly sensitive to sleep loss at ≈ P24. In rodents, this overlaps with critical periods (CP), which are developmental ages when the brain is exquisitely plastic and sensitive to changes in experience (Frank, 2011b, 2004). For example, sleep loss in mice during the visual CP impairs cortical plasticity necessary for normal development of binocular vision (Renouard et al., 2022, this issue; Zhou et al., 2020). Consistent with CPs in complex behavior, sleep disruption in early life leads to persistent alterations in social interactions and cognition in wild type rodents (voles) and in an autism mouse model (Jones et al., 2021, 2019; Lord et al., 2022; Milman et al., 2023). Collectively, these findings may explain why during the 3^rd^-4^th^ postnatal week (when many CPs occur in rodents) SD impacts PFC gene expression in ways not found at other ages.

For example, at P24, SD downregulates many more biological functions than it upregulates. These include growth factor binding factors (i.e. regulators of TGF-β, a cytokine with well-established roles in brain development (Meyers and Kessler, 2017)) and Wnt-signaling. (Figure 5, Supplementary table 4). Wnt-signaling is a pathway that is downregulated by SD at all ages (Figure 3), but this is most pronounced at P24 (Figure 6). Given the importance of both Wnt and TGF-β signaling for development and brain plasticity, the unique effect of SD in those pathways at P24 indicate this is an age when SD is particularly likely to disrupt both processes.

**Figure 6.**
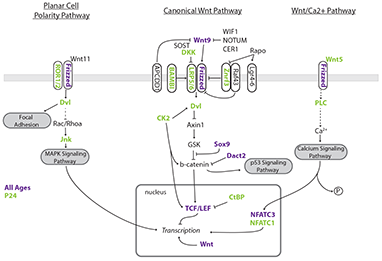
Genes related to in Wnt-signaling pathways are downregulated after SD at all ages but more extensively after SD in P24. Wnt-signaling pathways adapted from KEGG (mmu04310) and WikiPathways. Genes in blue are downregulated after SD in all ages. Genes colored in lime green are downregulated after SD only in P24.

### 4.4 SD-induced changes unique to the adult brain

There were several gene changes only observed in the adult brain (Fig.7), including the downregulation of genes involved in glycolysis/gluconeogenesis, pentose phosphate pathways, or lipid metabolism. This is consistent with the idea that different energy substrates are used by the awake and sleeping brain (Benington and Heller, 1995; Karnovsky, 1982; Karnovsky et al., 1983; Wisor et al., 2013). Indeed, processes related to elongation of fatty acids are significantly upregulated, more specifically Acetyl-CoA thioesterases (*ACOT,* Fig.7) and Hydroxyacyl-CoA dehydrogenase B (*Hadhb,* Fig.7). ACOTs hydrolyze Acetyl-CoA to fatty acids and CoASH (Hunt et al., 2006), while HADHB is involved in fatty acid beta oxidation (Dagher et al., 2021). Considering that brain energy demands are higher in early life (Kuzawa et al., 2014; Nehlig, 1997), it was surprising not to see similar changes in genes related to metabolism occur in P24-P30 mice. One possible explanation is that developing mice rely on different proportions of these substrates (Cremer, 1982; Nehlig, 1997; Prins, 2008). This is further supported by lower expression levels of cortical glucose transporters in younger animals (Vannucci, 1994), limiting glucose availability in the brain.

**Figure 7.**
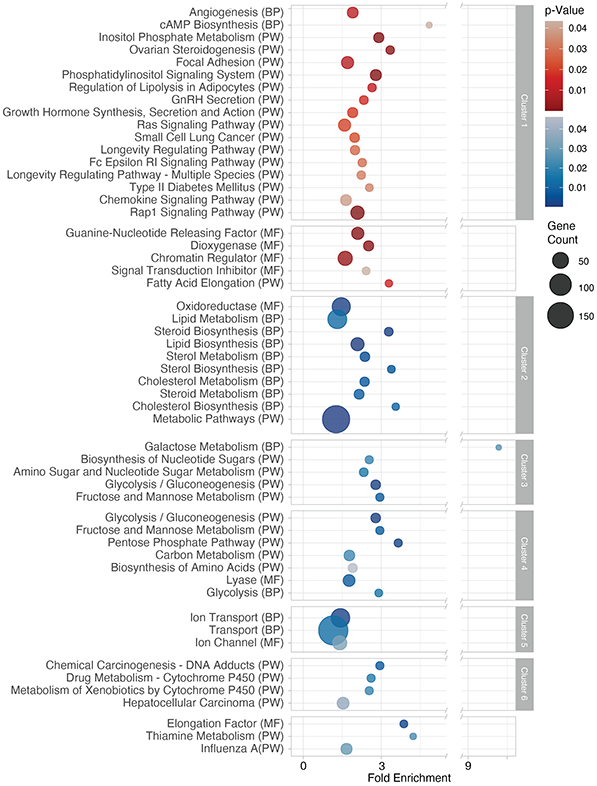
Differentially expressed genes after SD only in P90 mice are mostly related to cell adhesion and polarity and glucose and lipid metabolism. Bubbles show enriched functions of up (red) and down (blue) regulated genes. Bubble size represents number of genes per function. Modified Fisher’s exact p-value <0.05, minimum of 3 genes for each function or pathway. P-values are represented as a color gradient. Gray boxes outline function within each cluster with a similarity threshold 0.2 and Modified Fisher’s exact p-value<0.05. Enrichment scores for each cluster; cluster-1: 2.12, cluster-2: 2.57, cluster-3: 2.09, cluster-4: 2.09, cluster-5: 2.04, cluster-6: 1.75. MF, Uniprot keywords Molecular Function. BP, Uniprot keywords Biological Process. PW, KEGG pathways.

### 4.5 The paucity of molecular changes in P16 mice

A surprising finding was that SD had a small effect on PFC gene expression in P16 animals. We find only 85 differentially expressed genes compared to several thousand genes at older ages. This is even though rodents at similar ages exhibit very strong sleep drive and after SD show compensatory changes in sleep time, duration, or phasic activity (Frank, 2020). One possible explanation is that sub-cortical mechanisms necessary for behavioral sleep compensation are intact at these ages, but the cortical mechanisms are not (Blumberg et al., 2014). This may explain the absence of increases in NREM SWA after SD in rodents <P24. A second possible explanation is that sleep drive (as measured by NREM sleep SWA) may be primarily driven by the absence of NREM sleep (Achermann and Borbély, 2003, 1995, 1992). In adult animals, the amount of sleep lost with SD is mostly NREM sleep. This is because in adult mammals, NREM sleep represents ≈ 80% of total sleep time (Jones, 1991). However, this proportion is quite different in neonates, with NREM sleep representing between 30-50% of total sleep time (Frank, 2020). REM sleep is also very similar to waking, at least in terms of general cortical activation (Siegel, 2000). Therefore, the paucity of differential PFC gene expression after SD at P16 (when there is much more REM sleep) may reflect the fact that SD at these ages leads to less NREM sleep reduction, and/or that replacing REM sleep with wake incurs less sleep pressure.

### 4.6 Conclusions

In this study, we determined for the first time how SD impacts cortical gene expression across developmental ages where adult sleep homeostasis is absent (P16), emerging (P24-P30) and fully mature (P90). Overall, our main findings are that SD has dramatically different effects on PFC gene expression at different postnatal ages, and that the 3^rd^-4^th^ postnatal week (≈ P24) may be a period of vulnerability to disruption of plasticity pathways important for CPs. This suggests that sleep serves conserved functions across development, and different functions depending on postnatal age. Future investigations are now needed to characterize molecular changes across development that occur during sleep, particularly after SD. For example, in adult mice recovery sleep following SD induces separate molecular changes that provide clues to sleep function (Gerstner et al., 2016). Studies in developing mice may be equally revealing.

## Abbreviations

SD: sleep deprivation
HC: home cage
PFC: prefrontal cortex
P: postnatal day
ZT: zeitgeber time
EEG: electroencephalogram
NREM: non-rapid eye movement
REM: rapid eye movement
SWA: slow wave activity
CP: critical period
ELSD: early live sleep disruption

---

**Supplementary Figure 1** Upper Quartile (UQ) normalization did not separate the RNAseq dataset. **A)** Principal component analysis of UQ normalized RNAseq data detected age as a first principal component (PC1, 18.89%) but was unable to separate variability induced by SD. n=5-8 per age group and condition, open circles: home cage, closed circles: SD. **B)** Relative Log expression for each sample and condition using Upper Quantile normalization. Color code as in A. **C-F)** Volcano plots of normalized dataset per age group, **C)** P16 **D)** P24 **E)** P30 **F)** P90 Each gray dot represents an expressed gene, blue dots are genes significantly changed after SD. Absolute differentially expressed gene numbers are given in blue for each volcano plot on the left. For the P90 group, red dots represent known sleep dependent genes (supplementary Table 1). 71.6% of these genes were found as DEG with UQ normalization, FDR <0.05.

**Supplementary Table 1**

List of positive and negative genes used to establish RUVs normalization. Genes retrieved with RUVs in P90 are marked in bold.

**Supplementary Table 2**

Complete list of all differentially expressed genes (home cage vs. SD) per age after RUVs normalization, color coded in up (red) and down (blue) regulated genes. For the P90, control genes (supplementary Table 1) are marked in bold. FDR p<0.05.

**Supplementary Table 3**

List of genes extracted from the Venn diagram in Figure 2 used in downstream functional enrichment and clustering analysis. All overlapping genes of a Venn segment are in a separate tab. Up and down regulated genes are in the same tab, color coded (up red, down blue).

**Supplementary Table 4**

Enrichment analysis output from DAVID. Enriched functions or pathways for up (red) and downregulated (blue) genes and their respective clusters and enrichment scores. *indicates genes from functions used in the heatmaps.

## Acknowledgements

We thank Drs. Jonathan Wisor, Christopher Hayworth, and Ashley Ingiosi for constructive discussions and Dr. Kristan Singletary for proofreading.

This work was supported by K01NS104172 and R35GM147020 to LP and R01NS114780 to MGF.

