## Supplementary figures and images for "Ontogenesis of the molecular response to sleep loss"

### SupplementaryFigure_1

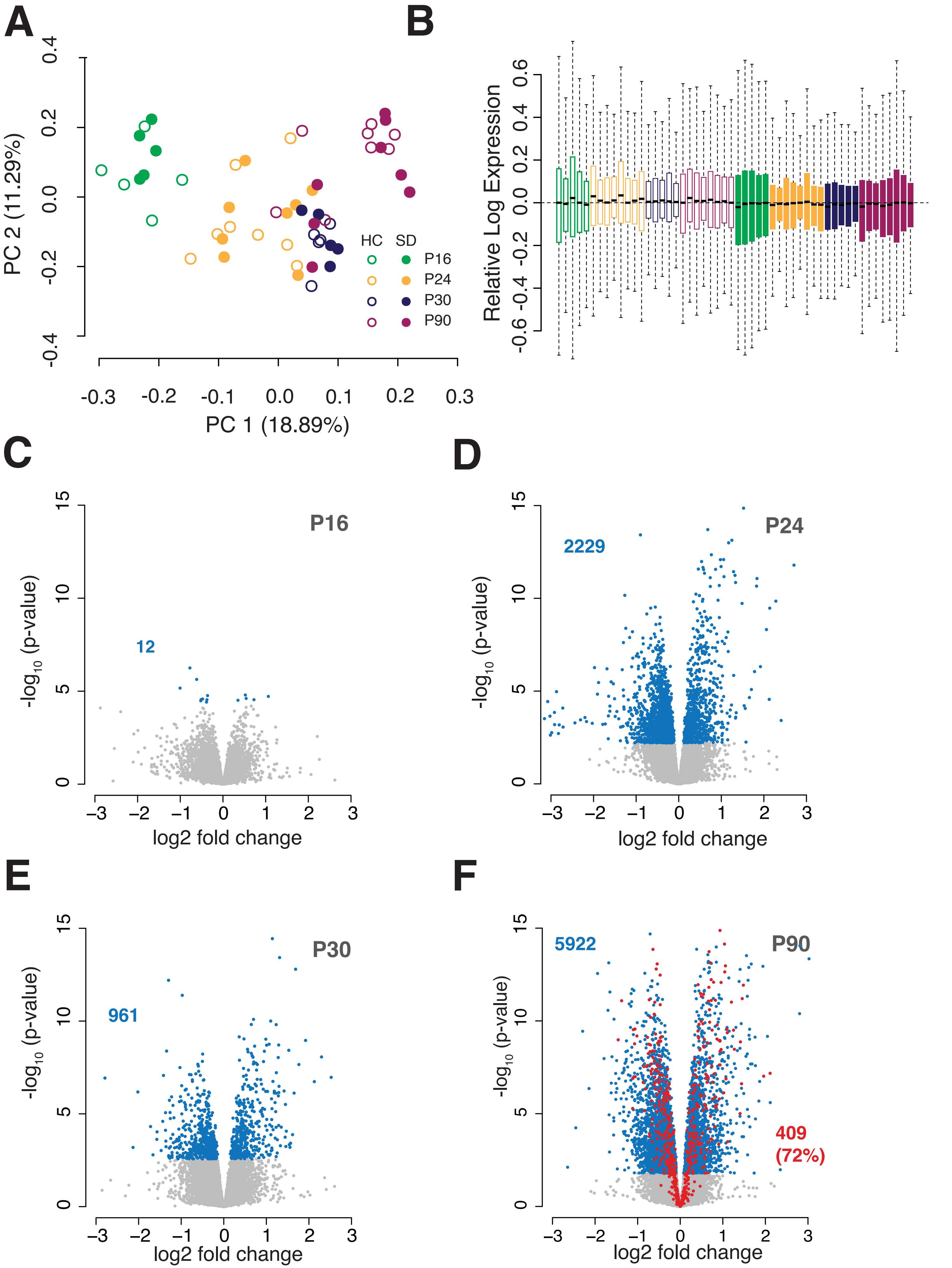
